# Anti-PF4 (heparin-independent)/PF4 complex induces allosteric activation of integrins αIIbβ3 and αvβ3, a potential mechanism of vaccine-induced thrombotic thrombocytopenia (VITT) and autoimmune diseases

**DOI:** 10.1101/2022.08.17.504306

**Authors:** Yoko K Takada, Chun-Yi Wu, Yoshikazu Takada

## Abstract

The classical immune-mediated heparin-induced thrombocytopenia (HIT) is induced by autoantibody against platelet-factor 4 (PF4)/heparin complex. Vaccine-induced thrombotic thrombocytopenia (VITT) and autoimmune HIT (aHIT) are induced by anti-PF4 in a heparin-independent manner. Activation of platelet integrin αIIbβ3 is a key event that leads to αIIbβ3 binding to fibrinogen and platelet aggregation, but is not involved in current models of HIT or VITT. Anti-PF4 (heparin-independent) is also detected in autoimmune diseases (e.g., SLE). However, the role of anti-PF4 in these diseases is unknown. Previous studies showed that several pro-inflammatory chemokines potently activated integrins by binding to the allosteric site (site 2). PF4 is known to be inhibitory since it inhibits angiogenesis and tumor growth. Here we describe that PF4 was predicted to bind to site 2 of αIIbβ3 by docking simulation, but did not activate it. However, PF4/anti-PF4 mAb (RTO, heparin-independent) complex potently activated it at biological concentrations of PF4 (<1 μg/ml), but anti-PF4/heparin (KKO) did not. This indicates that RTO changed the phenotype of PF4. We generated PF4 mutants defective in site 2 binding to integrin by introducing mutations in the predicted site 2 binding site of PF4. A PF4 mutant/RTO complex was defective in activating integrins. Furthermore, this PF4 mutant acted as an antagonist of PF4/RTO-induced integrin activation. We obtained similar results with vascular integrin αvβ3. We propose that a potential mechanism, in which PF4/RTO complex binds to site 2 and activates integrins and triggers thrombocytopenia or autoimmune diseases. The inhibitory PF4 mutant may have potential as a therapeutic.

## Introduction

It has been reported that vaccination for SARS-CoV-2 may result in a vaccine-induced catastrophic thrombotic thrombocytopenia (VITT) disorder. This disorder presents as extensive thrombosis in atypical sites, primarily in the cerebral venous, alongside thrombocytopenia and the production of autoantibody against platelet-factor 4 (PF4, chemokine CXCL4). PF4 is one of the most abundant proteins in platelet granules and PF4 is present at > 1 μg/ml concentrations in plasma. This rare adverse effect extremely resembles the clinical presentation of the classical immune-mediated heparin-induced thrombocytopenia (HIT) disorder, which is induced by anti-PF4/heparin complex and occurs following exposure to heparin (3-10). VITT is also very similar to autoimmune HIT (aHIT), which is induced by anti-PF4 but none of these patients had been pre-exposed to heparin before disease onset.

Anti-PF4 autoantibodies have also been detected in several autoimmune diseases (e.g., SLE, Systemic sclerosis, and RA)(11-13). It is unclear how anti-PF4 induces thrombocytopenia. A current model of thrombotic thrombocytopenia suggests that (1) anti-PF4 binds to PF4 and induces PF4 clustering, (2) the complex binds to platelets by binding to the FcRγIIA receptor and proteoglycans of platelets, leading to (3) platelet activation and aggregation (10).

Integrins are a superfamily of αβ heterodimers that were originally identified as receptors for extracellular matrix proteins (14). PF4 is known to bind to αvβ3 (15) and Mac-1 (16). It is unclear if PF4 binds to αIIbβ3 and activates it. We previously discovered that the chemokine domain of pro-inflammatory chemokine CX3CL1 is a ligand for integrins αvβ3 and α4β1 and bound to the classical ligand-binding site of integrins (site 1) (17). We showed that CX3CL1 activated soluble integrin αvβ3 in cell-free conditions in ELISA-type activation assay (18). In this assay, we coated the wells of 96-well plate with specific integrin ligands and incubated with soluble integrins in the presence of CX3CL1 in 1 mM Ca^2+^ to keep integrin inactive. We detected the increase in integrin binding to immobilized ligand γC399tr, a fibrinogen fragment, indicating that soluble integrins were activated by CX3CL1. CX3CL1 binds to the allosteric site (site 2) in integrin headpiece, which is distinct from the classical ligand-binding site (site 1) (18). Site 2 is located on the opposite side of site 1 in the integrin headpiece **(Fig. 1)**. Site 2 was identified by docking simulation of the interaction between the closed/inactive integrin αvβ3 (1JV2.pdb) and CX3CL1 (18). Another pro-inflammatory chemokines SDF-1 (CXCL12) activated integrins αvβ3, α4β1, and α5β1 by binding to site 2 (19). 25-Hydroxycholesterol, a mediator of inflammatory signals in innate immunity, was shown to bind to integrin site 2 and induce integrin activation and inflammatory signaling, leading to over-production of inflammatory cytokines (e.g., IL-6 and TNFα) in monocytes (20). It has thus been proposed that site 2 plays a critical role in inflammation.

**Fig. 1.**
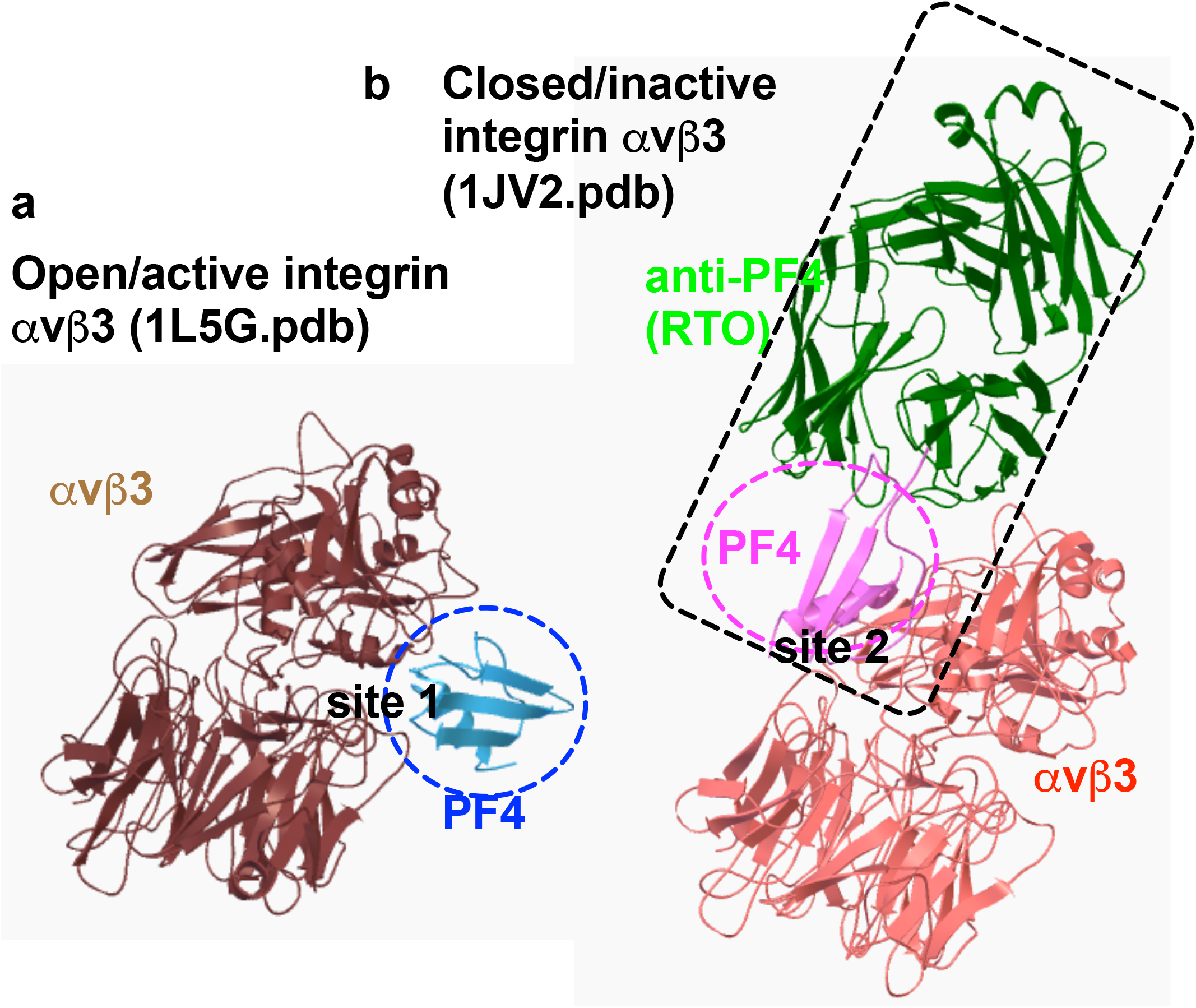
Docking models of anti-PF4/PF4-integrin interaction. <u>(a) PF4 binding to the classical ligand-binding site (site 1) of active αvβ3 (1L5G.pdb).</u> 3D structure of αvβ3 was used because active and inactive 3D structures are known. Autodock3 was used for docking simulation. The simulation predicts that PF4 binds to site 1 (docking energy - 23.4 kcal/mol). <u>(b) PF4 binding to the allosteric site (site 2) of inactive αvβ3 (1 JV2.pdb).</u> Docking energy −21.2 kcal/mol. When anti-PF4 (RTO)/PF4 complex structure (4RAU.pdb) was superposed, PF4/anti-PF4 is predicted to bind to αvβ3 (site 2) without steric hindrance. We hypothesize that PF4 binds to the site 2 of inactive integrins but does not induce activation at biological concentrations of PF4 (Figs. 3 and 4). The anti-PF4/PF4 complex induces integrin activation by changing conformation of PF4. Anti-PF4 is detected in thrombocytopenia and other autoimmune diseases, and this activation by anti-PF4/PF4 may be potentially involved in the pathogenesis of diseases. Docking simulation with the 3D structure of αIIbβ3 was similar.

We thus hypothesized that anti-PF4 regulates PF4-induced activation of integrins. In the present study, we showed that PF4 did not activate integrins by itself although PF4 is predicted to bind to site 2. In contrast, anti-PF4/PF4 complex potently activated integrins αIIbβ3 and αvβ3. We generated PF4 mutants defective in site 2 binding. A PF4 mutant/anti-PF4 complex was defective in activating integrins αIIbβ3 and αvβ3, and acted as an antagonist of PF4/anti-PF4-induced integrin activation. These findings suggest that anti-PF4/PF4 requires to bind to site 2 for activating integrins. We propose that anti-PF4/PF4 complex induces integrin αIIbβ3 activation, and induce subsequent αIIbβ3-fibrinogen bridge, leading to platelet aggregation. Also, anti-PF4 may be involved in the pathogenesis of autoimmune diseases by activation of αvβ3 and other integrins in non-platelet cells (e.g., monocytes). The PF4 mutant defective in binding to site 2 is a potential antagonist of anti-PF4-induced VITT and autoimmune diseases.

## Results

### PF4 specifically binds to site 1 of integrins αIIbβ3 and αvβ3

Although PF4 is known to bind to integrins αvβ3, the specifics of interaction are unclear. To predict how PF4 binds to integrin, we performed docking simulation of interaction between PF4 (1RHP.pdb) and integrin αvβ3 using autodock3. The 3D structure of αvβ3 was used since active and inactive conformers are well defined. In our docking studies, PF4 is predicted to bind to site 1 (docking energy −24.3 kcal/mol) of active conformer of integrin αvβ3 (open headpiece/active, 1L5G.pdb)(Fig. 1a). PF4 bound to soluble αIIbβ3 and αvβ3, which are activated by 1 mM Mn^2+^ in ELISA type binding assays in cell-free conditions (Fig. 2a), consistent with the prediction. We found that the disintegrin domain of ADAM15, which is known to bind to integrins αvβ3 (21) and αIIbβ3 (22), inhibited the binding of PF4 to integrins, but control GST did not (Fig. 2b and 2c). This indicates that the PF4 binding to these integrins is specific.

**Fig. 2.**
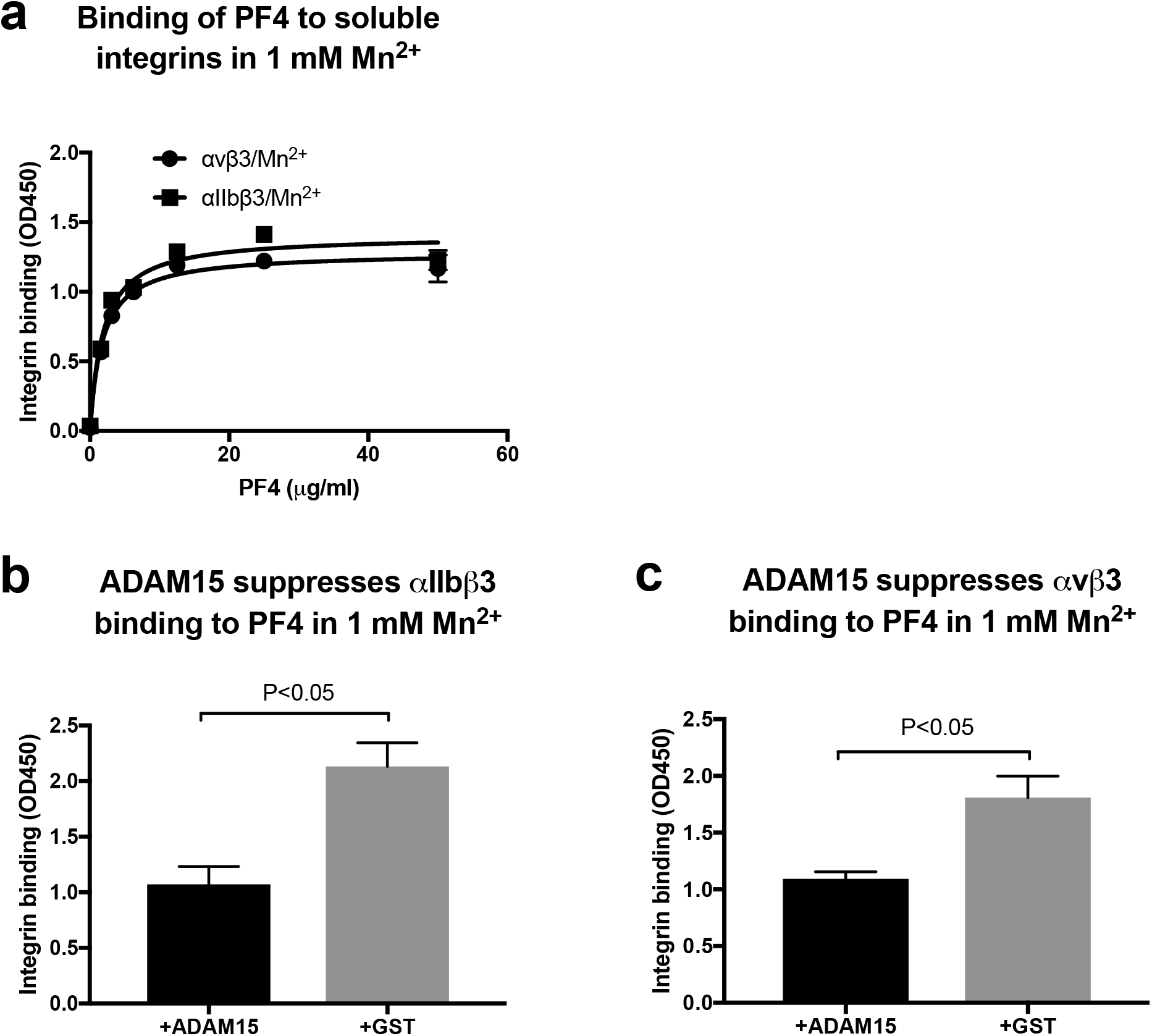
PF4 specifically binds to soluble αIIbβ3 and αvβ3 in ELISA-type binding assays in 1 mM Mn^2+^. a) PF4 binds to soluble integrins in 1 mM Mn^2+^ in ELISA-type <u>binding assays.</u> PF4 was immobilized to wells of 96-well microtiter plate and incubated with soluble αIIbβ3 or αvβ3 (1 μg/ml) in Tyrode-HEPES buffer with 1 mM Mn^2+^ (to activate integrins) for 1 hr at room temperature and bound integrins were quantified using anti-β3 (mAb AV10) and HRP-conjugated anti-mouse IgG. The data show that PF4 binds to these integrins at Kd <1 μg/ml. (n=3) <u>(b and c) The binding of PF4 to integrins was suppressed by the distintegrin domain of ADAM15 fused to GST (ADAM15 disintegrin), but not by control GST.</u> To establish the specificity of PF4 binding to soluble integrins αIIbβ3 (b) and αvβ3 (c), if ADAM15 disintegrin, which is known to bind to integrins αIIbβ3 (22) and αvβ3 (21), suppress the binding. ADAM15 disintegrin (100 μg/ml) suppressed the integrin binding to immobilized PF4 (12.5 μg/ml). This indicates that the binding of soluble integrins to PF4 is specific.

### PF4 is predicted to bind to site 2, but did not activate integrins

We studied if PF4 binds to the allosteric site of integrins (site 2) and activates integrins, as in pro-inflammatory chemokines CX3CL1 and CXCL12 (18, 19). Docking simulation of interaction between PF4 (1RHP.pdb) and inactive conformer of integrin αvβ3 (closed headpiece/inactive 1JV2.pdb) predicted that PF4 binds to site 2 (docking energy 21 kcal/mol)(Fig. 1b), predicting that PF4 allosterically activate integrins as in CX3CL1 or CXCL12. However, PF4 by itself did not activate αIIbβ3 or αvβ3 at biological concentrations (<1 μg/ml) (Figs. 3a and 4a). We thus hypothesized that PF4 signaling upon binding to integrins can be modified by anti-PF4.

**Fig. 3.**
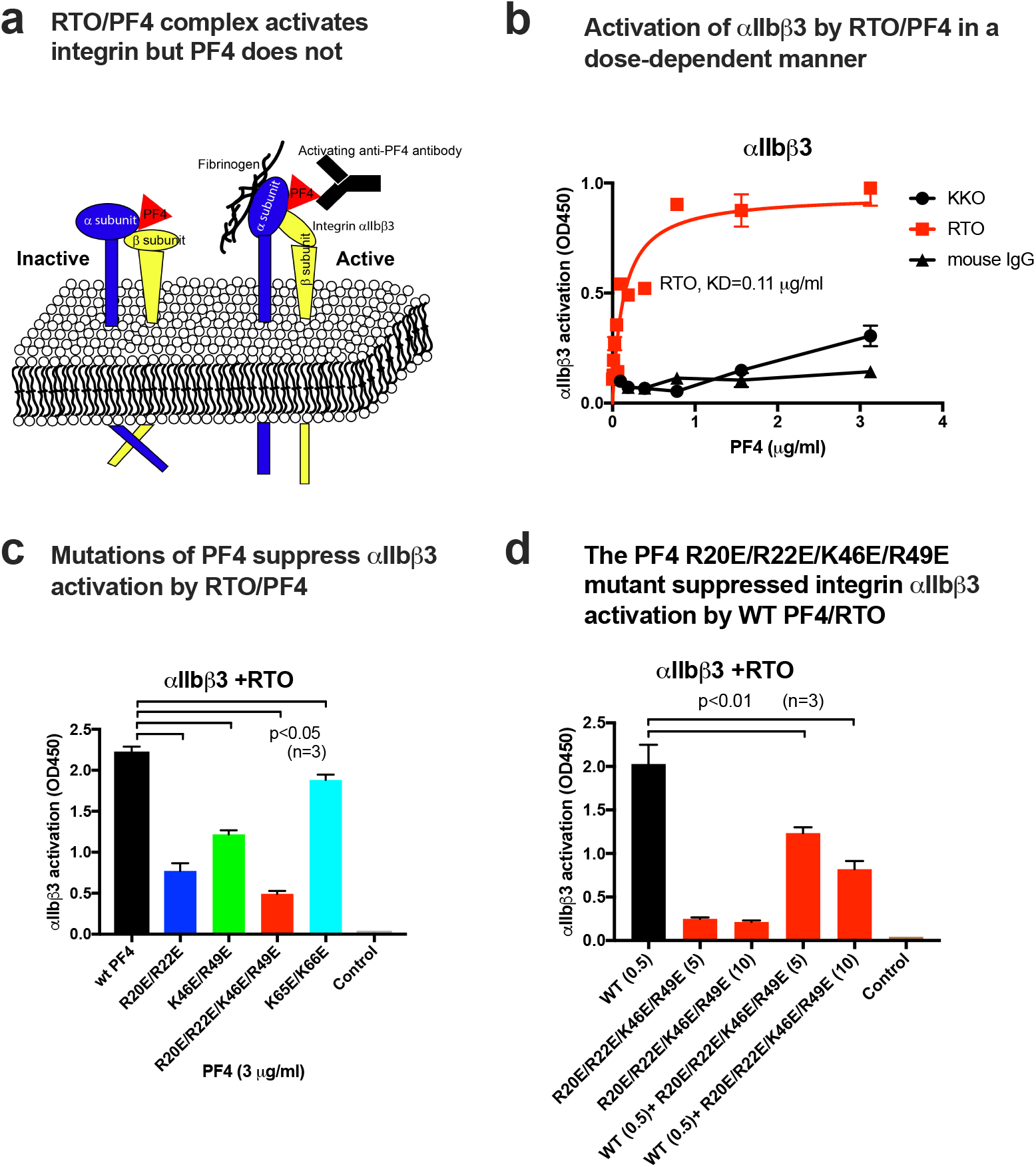
Anti-PF4/PF4 complex potently activated integrin αIIbβ3, but PF4 itself did not. a. <u>A model of PF4-induced integrin activation</u>. b. <u>Anti-PF4 (RTO) markedly</u> enhances PF4-induced activation of soluble αIIbβ3 in 1 mM Ca^2+^ in ELISA-type <u>activation assays.</u> The fibrinogen fragments γC390-411 (a specific ligand for αIIbβ3) was immobilized to wells of 96-well microtiter plate. Wells were incubated with soluble αIIbβ3 (1 μg/ml) in Tyrode-HEPES buffer with PF4 in 1 mM Ca^2+^ (to keep integrins inactive) for 1 hr at room temperature. Anti-PF4 (RTO or KKO, 10 μg/ml) were added (without heparin). After washing, bound integrins were quantified using anti-β3 (mAb AV10) and HRP-conjugated anti-mouse IgG. After washing, bound integrins were quantified using anti-β3 (mAb AV10) and HRP-conjugated anti-mouse IgG. Mouse IgG (10 μg/ml) and no antibody were used as negative controls. (n=3) The data show that PF4 itself did not activate αIIbβ3 at <1 μg/ml, but the RTO/PF4 complex did. <u>c. Point mutations in the predicted integrin-binding site (site 2) of PF4 suppressed activation of αIIbβ3 by the anti-PF4/PF4 complex.</u> The PF4 mutant (R20E/R22E/K46E/R49E) most effectively suppressed integrin activation by anti-PF4/PF4 complex. d. The R20E/R22E/K46E/R49E mutant suppressed integrin activation by the anti-PF4/PF4 complex. WT PF4 (0.5 μg/ml) and excess PF4 mutant (5 or 10 μg/ml) were used. The data indicate that the R20E/R22E/K46E/R49E mutant acted as an antagonist.

**Fig. 4.**
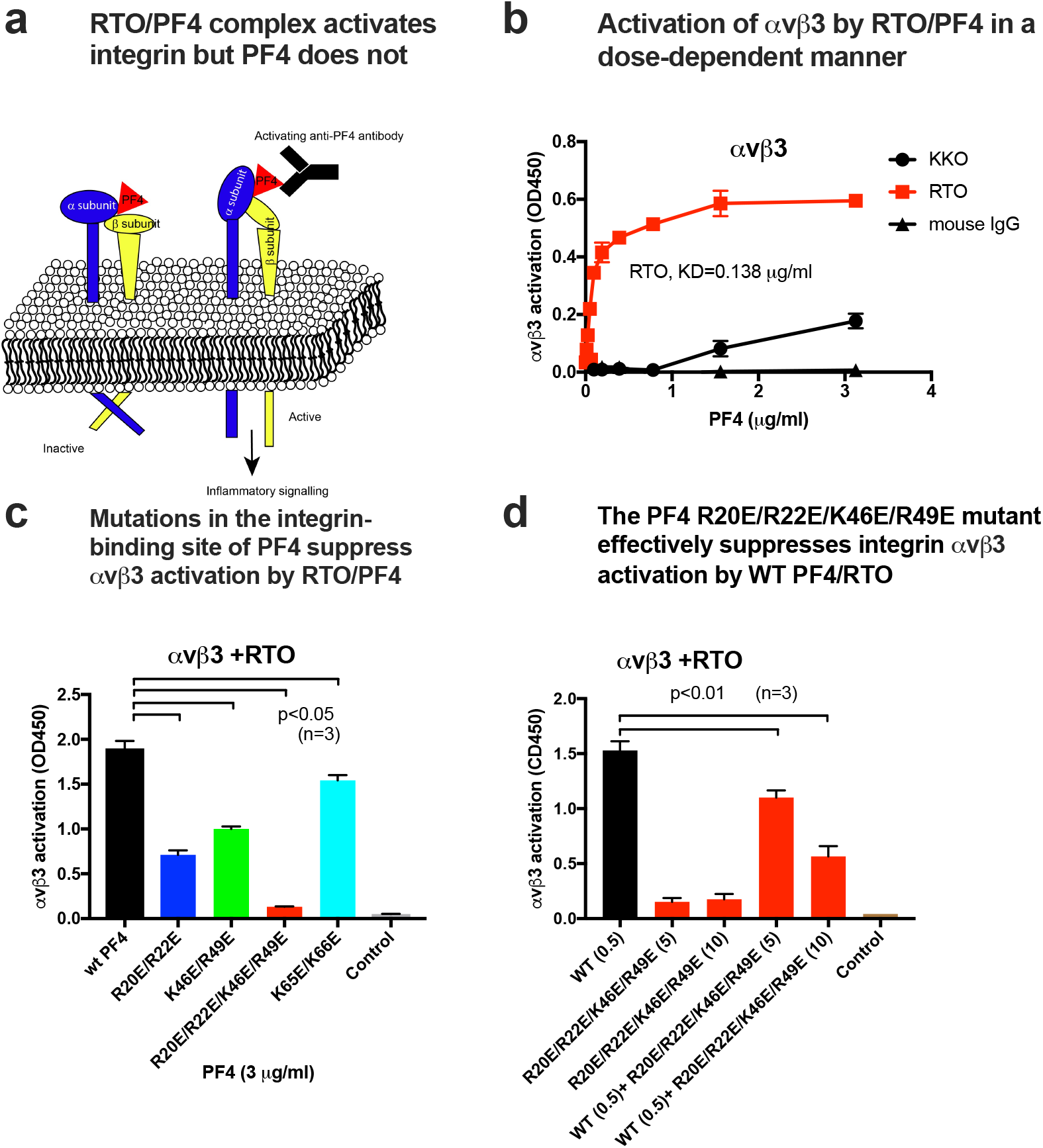
Anti-PF4/PF4 complex potently activated integrin αvβ3, but PF4 itself did not. a. <u>A model of PF4-induced integrin activation</u>. b. <u>Anti-PF4 (RTO) markedly</u> enhances PF4-induced activation of soluble αvβ3 in 1 mM Ca^2+^ in ELISA-type activation <u>assays.</u> The fibrinogen fragments γC390-411 (a specific ligand for <u>αvβ3</u>) was immobilized to wells of 96-well microtiter plate. Wells were incubated with soluble <u>αvβ3</u> (1 μg/ml) in Tyrode-HEPES buffer with PF4 in 1 mM Ca^2+^ (to keep integrins inactive) for 1 hr at room temperature. Anti-PF4 (RTO or KKO, 10 μg/ml) were added (without heparin). After washing, bound integrins were quantified using anti-β3 (mAb AV10) and HRP-conjugated anti-mouse IgG. After washing, bound integrins were quantified using anti-β3 (mAb AV10) and HRP-conjugated anti-mouse IgG. Mouse IgG (10 μg/ml) and no antibody were used as negative controls. (n=3) The data show that PF4 itself did not activate αvβ3 at <1 μg/ml, but the RTO/PF4 complex did. <u>c. Point mutations in the predicted integrin-binding site (site 2) of PF4 suppressed activation of αvβ3 by the anti-PF4/PF4 complex.</u> The PF4 mutant (R20E/R22E/K46E/R49E) most effectively suppressed integrin activation by anti-PF4/PF4 complex. d. The R20E/R22E/K46E/R49E mutant suppressed integrin activation by the anti-PF4/PF4 complex. WT PF4 (0.5 μg/ml) and excess PF4 mutant (5 or 10 μg/ml) were used. The data indicate that the R20E/R22E/K46E/R49E mutant acted as an antagonist.

### PF4/anti-PF4 (RTO) complex potently activates integrins at physiological PF4 concentrations

A murine mAb KKO to human (h) PF4/heparin complexes is known to bind specifically to hPF4/heparin complexes and induces HIT (pathogenic mAb) (23). Murine anti-hPF4 mAb RTO does not require heparin for binding to PF4 and does not induce HIT (non-pathogenic) (23). We studied if RTO and KKO influence the ability of PF4 to induce integrin activation. We expected that pathogenic KKO will activate αIIbβ3 and αvβ3 by binding to PF4 and non-pathogenic RTO will not. Unexpectedly, the RTO (at 10 μg/ml)/PF4 complex markedly activated integrins but KKO/PF4 complex did not in a heparin-independent manner in ELISA-type activation assays in 1 mM Ca^2+^. KKO/PF4 complex did not (Fig. 3b and 4b). This is the first clue that anti-PF4/PF4 complex is potentially involved in integrin αIIbβ3 activation, a trigger of platelet aggregation. Since heparin-independent anti-PF4 has been detected in autoimmune diseases and the levels of anti-PF4 correlate with disease progression (11, 13), activation of vascular αvβ3 by anti-PF4/PF4 complex may be involved in the pathogenesis of autoimmune diseases. RTO can be used as a model of heparin-independent anti-PF4 detected in VITT, aHIT and autoimmune diseases.

### PF4 mutant defective in site 2 binding is defective in integrin activation and acted as an antagonist for PF4/RTO-induced integrin activation

Table 1 shows amino acid residues in PF4 that are involved in site 2 binding. We superposed the PF4/RTO complex (1RHP.pdb) and PF4/αvβ3 complex. It is predicted that RTO, PF4 and integrin can co-exist without steric hindrance. The predicted RTO-binding site and site 2-binding site in PF4 are distinct. To determine if PF4/RTO complex activates integrins by binding to site 2, we developed PF4 mutants that are defective in site 2 binding by introducing mutations in the site 2-binding interface in PF4 predicted by docking simulation (Fig. 1). Arg20, Arg22, Lys46, Arg49, Lys65, and Lys66 in the integrin-binding interface of PF4 were mutated to Glu. The R20E/R22E and K46E/R49E mutations showed reduced ability to mediate RTO/PF4-induced integrin activation (Fig. 3c and 4c). The combined PF4 mutant (R20E/R22E/K46E/R49E) most effectively reduced RTO/PF4-induced integrin activation (Fig. 3c and 4c). Notably, this mutant suppressed integrin activation induced by PF4/RTO complex in a dose-dependent manner (dominant-negative effect) (Fig. 3d and 4d). This PF4 mutant binds to anti-PF4 (heparin-independent) but cannot induce integrin activation since it can not bind to site 2. The PF4 mutant competes with wild type PF4 complex for binding to anti-PF4.

**Table 1.** Amino acid residues involved in PF4-integrin αvβ3 interaction predicted by docking simulation

| PF4 (1RHP.pdb) | $\alpha v$ (1JV2.pdb) | $\beta 3$ (1JV2.pdb) |
| --- | --- | --- |
| Thr15, Thr16, Ser17, Gln18, Val19, **Arg20**, Pro21, **Arg22**, His23, Thr25, **Lys46**, Asn47, **Arg49**, Asp54, Leu55, Gln56, Ala57, Pro58, Leu59, **Lys62**, **Lys65**, **Lys66**, Leu67, Glu69, Ser70 | Glu15, Gly16, Ser17, Tyr18, Phe19, Lys42, Asn44, Ile50, Val51, Glu52, Trp93, Ala396, Ala397, Arg398, Ser399, Met400, Phe427, Gly428, Val429, Asp430 | Pro160, Met165, Lys235, Gly264, Ile265, Val266, Gln267, Asp270, Gln272, Cys273, His274, Val275, Gly276, Ser277, Asp278, His280, Tyr281, Ser282, Ala283, Thr285, Thr286, Met287 |

Amino acid residues within 0.6 nm between PF4 and $\alpha v\beta 3$ were selected using pdb viewer (version 4.1). Amino acid residues in PF4 selected for mutagenesis are shown in bold.

## Discussion

The present study establishes that RTO/PF4 complex potently activated integrins by binding to site 2, although PF4 by itself did not activate integrins at physiological concentrations of PF4. Since the RTO/PF4 mutant defective in binding to site 2 did not activate integrins and acted as an antagonist of anti-PF4/PF4-induced integrin activation, PF4 binding to site 2 is critical for RTO/PF4-induced integrin activation. It is likely that RTO binding to PF4 changed the phenotype of PF4 and activated integrins upon binding to site 2. This phenomenon mimics the anti-PF4-induced aHIT and VITT, and potentially connects thrombocytopenia, platelet integrin αIIbβ3, and anti-PF4. It is interesting that KKO (heparin-dependent anti-human PF4 induces HIT, but RTO (heparin-independent anti-human PF4) does not (4). One possibility is that heparin-independent and dependent TT are induced by totally different mechanisms. The PF4 mutant defective in site 2 binding may suppress thrombosis by blocking anti-PF4-induced integrin αIIbβ3 activation. Therefore, RTO is a model of heparin-independent autoantibodies to PF4 that induce thrombocytopenia. Our preliminary studies predict that anti-PF4/PF4-induced αIIbβ3 activation causes thrombocytopenia. We should urgently address this hypothesis. We propose that anti-PF4 like RTO can change the phenotype of PF4 and anti-PF4/PF4 complex can activate integrins by binding to site 2. The PF4 mutant defective in site 2 binding has potential as an antagonist for allosteric integrin activation. The ELISA-type activation of integrins by anti-PF4/PF4 complex can be potentially useful to detect heparin-independent anti-PF4 in patients’ blood.

Also, RTO/PF4 complex can activate vascular integrin αvβ3 in an allosteric manner. Since anti-PF4 is detected in several autoimmune diseases as well, and PF4 does not activate αvβ3 (and perhaps other integrins), anti-PF4 stimulate integrin activation in cell types (e.g., monocytes) other than platelets. The levels of heparin-independent anti-PF4 is known to correlate with disease activity index in SLE patients (11). Therefore, it is likely that the same PF4 mutant may block vascular inflammation induced by anti-PF4 in autoimmune diseases.

## Experimental procedures

### Fibrinogen γ-chain C-terminal residues 390-411, a specific ligand for αIIbβ3 fused to GST

cDNA encoding (6 His tag and Fibrinogen γ-chain C-terminal residues 390-411) [HHHHHH]NRLTIGEGQQHHLGGAKQAGDV] was conjugated with the C-terminus of GST (designated γC390-411) in pGEXT2 vector (BamHI/EcoRI site). The protein was synthesized in E. coli BL21 and purified using glutathione affinity chromatography. <u>The fibrinogen γ-chain C-terminal domain (γC399tr, residues 151-399)</u>, a specific ligand for αvβ3 has been previously described (24). The disintegrin domain of ADAM15 fused to GST (ADMA15 disintegrin) and parent GST were synthesized as previously described (21).

#### PF4

The cDNA encoding PF4 was synthesized and subcloned into the BamHI/EcoRI site of pET28a vector. Protein synthesis was induced by IPTG in E. coli BL21 and protein was synthesized as insoluble inclusion bodies and purified in Ni-NTA affinity chromatography under denaturing conditions and renatured as described (17).

Anti-PF4 antibodies RTO and KKO were obtained from Invitrogen.

### ELISA-type integrin activation assays (19)

Wells of 96-well microtiter plates were coated with γC390-411 (a specific ligand for αIIbβ3) or γC399tr (a specific ligand for αvβ3). Remaining protein binding sites were blocked with BSA. Soluble recombinant αIIbβ3 or αvβ3 (AgroBio, 1 μg/ml) was pre-incubated with PF4 or anti-PF4/PF4 for 10 min at room temperature and was added to the wells and incubated in HEPES-Tyrodes buffer with 1 mM CaCl2 for 1 h at room temperature. After unbound αIIbβ3 or αvβ3 was removed by rinsing the wells with binding buffer, bound αIIbβ3 or αvβ3 was measured using anti-integrin β3 mAb (AV-10) followed by HRP-conjugated goat anti-mouse IgG and peroxidase substrates.

### ELISA-type integrin binding assays (17)

Wells of 96-well microtiter plates were coated with PF4. Remaining protein binding sites were blocked by incubating with BSA. After washing with PBS, soluble recombinant αIIbβ3 or αvβ3 (1 μg/ml) was added to the wells and incubated in HEPES-Tyrodes buffer with 1 mM MnCl2 for 1 h at room temperature. Bound αIIbβ3 or αvβ3 was measured using anti-integrin β3 mAb (AV-10) followed by HRP-conjugated goat anti-mouse IgG and peroxidase substrates.

#### Docking simulation

Docking simulation of interaction between PF4 and integrin αvβ3 (open headpiece form 1L5G, or closed headpiece form, PDB code 1JV2) was performed using AutoDock3 as described previously (1). We used the headpiece (residues 1–438 of αv and residues 55–432 of β3) of αvβ3. Cations were not present in integrins during docking simulation. The ligand is presently compiled to a maximum size of 1024 atoms. Atomic solvation parameters and fractional volumes were assigned to the protein atoms by using the AddSol utility, and grid maps were calculated by using AutoGrid utility in AutoDock 3.05. A grid map with 127 × 127 × 127 points and a grid point spacing of 0.603 Å included the headpiece of αvβ3. Kollman ‘united-atom’ charges were used. AutoDock 3.05 uses a Lamarckian genetic algorithm (LGA) that couples a typical Darwinian genetic algorithm for global searching with the Solis and Wets algorithm for local searching. The LGA parameters were defined as follows: the initial population of random individuals had a size of 50 individuals; each docking was terminated with a maximum number of 1 × 10^6^ energy evaluations or a maximum number of 27 000 generations, whichever came first; mutation and crossover rates were set at 0.02 and 0.80, respectively. An elitism value of 1 was applied, which ensured that the top-ranked individual in the population always survived into the next generation. A maximum of 300 iterations per local search were used. The probability of performing a local search on an individual was 0.06, whereas the maximum number of consecutive successes or failures before doubling or halving the search step size was 4.

#### Other methods

Treatment differences were tested using ANOVA and a Tukey multiple comparison test to control the global type I error using Prism 7 (Graphpad Software).

